# A distinct innate immune signature marks progression from mild to severe COVID-19

**DOI:** 10.1101/2020.08.04.236315

**Authors:** Stéphane Chevrier, Yves Zurbuchen, Carlo Cervia, Sarah Adamo, Miro E. Raeber, Natalie de Souza, Sujana Sivapatham, Andrea Jacobs, Esther Bächli, Alain Rudiger, Melina Stüssi-Helbling, Lars C. Huber, Dominik J. Schaer, Jakob Nilsson, Onur Boyman, Bernd Bodenmiller

## Abstract

Coronavirus disease 2019 (COVID-19) manifests with a range of severities, but immune signatures of mild and severe disease are still not fully understood. Excessive inflammation has been postulated to be a major factor in the pathogenesis of severe COVID-19 and innate immune mechanisms are likely to be central in the inflammatory response. We used 40-plex mass cytometry and targeted serum proteomics to profile innate immune cell populations from peripheral blood of patients with mild or severe COVID-19 and healthy controls. Sampling at different stages of COVID-19 allowed us to reconstruct a pseudo-temporal trajectory of the innate immune response. Despite the expected patient heterogeneity, we identified consistent changes during the course of the infection. A rapid and early surge of CD169^+^ monocytes associated with an IFNγ^+^MCP-2^+^ signature quickly followed symptom onset; at symptom onset, patients with mild and severe COVID-19 had a similar signature, but over the course of the disease, the differences between patients with mild and severe disease increased. Later in the disease course, we observed a more pronounced re-appearance of intermediate/non-classical monocytes and mounting systemic CCL3 and CCL4 levels in patients with severe disease. Our data provide new insights into the dynamic nature of the early inflammatory response to severe acute respiratory syndrome coronavirus 2 (SARS-CoV-2) infection and identifies sustained pathological innate immune responses as a likely key mechanism in severe COVID-19, further supporting investigation of targeted anti-inflammatory interventions in severe COVID-19.

## Introduction

Coronavirus disease 2019 (COVID-19) was first identified in December 2019 in Wuhan, China (Zhu *et al.*, 2020). The disease developed into a global pandemic with over 15 million confirmed cases and over 600,000 confirmed deaths as of July 24th 2020 (Dong, Du and Gardner, 2020). The clinical presentation of COVID-19 can vary from asymptomatic or mild cases to an acute respiratory distress syndrome (ARDS) requiring mechanical ventilation (Wu and McGoogan, 2020). About 5% of those clinically diagnosed with COVID-19 develop ARDS; these patients generally experience a sudden deterioration after around 1 week of symptom onset (Wiersinga *et al.*, 2020).

SARS-CoV-2, a positive-sense, single-stranded RNA virus, has been identified as the causative pathogen of COVID-19. This virus shows a tropism for cells that express the angiotensin-converting enzyme 2 (ACE2), which serves as an entry receptor for SARS-CoV-2 into cells of the respiratory tract, kidneys, liver, heart, brain, and blood vessels (Puelles *et al.*, 2020). Upon infection of epithelial cells, pattern recognition receptors that sense viral RNA, such as TLR7 and 8, initiate interferon (IFN) production and innate immune cell recruitment, triggering an inflammatory response that, in COVID-19, has also been linked to inflammasome activation (Iwasaki and Medzhitov, 2015; Yap, Moriyama and Iwasaki, 2020).

Early data from China indicated that patients with severe disease mount a strong inflammatory response as shown by increased levels of proinflammatory cytokines, such as tumor necrosis factor (TNF), monocyte chemoattractant protein 1 (MCP-1, also known as CCL2), and macrophage inflammatory protein 1α (MIP-1α, also known as CCL3) (Huang *et al.*, 2020). These data have been confirmed in other studies, which also revealed a distinct cytokine response with activated IL-1 and IL-6 pathways and chemokine enriched signatures (Blanco-Melo *et al.*, 2020; Veerdonk *et al.*, 2020). The type I IFN response during SARS and SARS-CoV-2 infection has gained particular attention; a suppressed or delayed type I IFN response may be associated with a severe disease course, potentially through the recruitment of proinflammatory monocytes to lungs even though the data are currently not conclusive (Channappanavar *et al.*, 2016; Hadjadj *et al.*, 2020; Lee *et al.*, 2020; Park and Iwasaki, 2020).

Myeloid cells have been implicated in the pathophysiology of COVID-19 by contributing to local tissue damage and as potential producers of the cytokines that lead to the systemic inflammatory state seen in patients with severe disease (McKechnie and Blish, 2020; Merad and Martin, 2020; Vabret *et al.*, 2020). Several studies have shown distinct changes within the monocytic compartment in patients with severe COVID-19 that were similar to an immune paralysis phenotype described in sepsis. Thus, monocytes were shown to downregulate HLA-DR while retaining the ability to secrete proinflammatory cytokines (Giamarellos-Bourboulis *et al.*, 2020). A recent single-cell RNA-sequencing study of bronchioalveolar lavage fluid showed changes in the local myeloid environment toward a proinflammatory, peripheral monocyte-derived phenotype and a depletion of alveolar macrophages in severe COVID-19 patients (Liao *et al.*, 2020). Using the same approach, others demonstrated a phenotypic shift of the CD14^+^ population and a depletion of CD16^+^ cells in the myeloid compartment in peripheral blood of COVID-19 patients compared to healthy individuals (Wilk *et al.*, 2020).

These data suggest a distinct role of the myeloid compartment in the pathogenesis of COVID-19; however, these studies have relied on relatively small sample sizes lacking patients with mild disease, and data regarding the cellular innate immune response and the underlying cytokine and chemokine network, at high phenotypic- and temporal resolution, is still missing. Here we describe an in-depth characterization of the myeloid compartments of 66 patients with mild to severe COVID-19 and 22 healthy controls by using 40-parameter mass cytometry and targeted proteomics of serum samples. Using this systems approach, we could reconstruct phenotypic changes arising throughout the course of the disease. At an early stage, the innate immune response was relatively similar in patients with mild and severe disease, characterized by an increase in CD169^+^ monocytes, which correlated with a strong pro-inflammatory cytokine signature. At a later stage, while patients with mild COVID-19 showed a normalization of their innate immune signature, patients with severe disease exhibited an ongoing inflammatory state dominated by a chemokine enriched signature and a higher frequency of CD16+ monocytes.

## Results

### Clinicopathological assessment of mild versus severe COVID-19 patients

To better understand the role of the myeloid compartment in the pathophysiology of COVID-19, we established a multicenter cohort of 66 COVID-19 patients. We have previously described the SARS-CoV-2-specific antibody response in a subset of this cohort (Cervia *et al.*, 2020). According to the WHO definition (World Health Organization, 2020) 28 of these patients experienced a mild disease course with either mild illness or mild pneumonia, whereas 38 experienced a severe disease course consisting of severe pneumonia or acute respiratory distress syndrome (ARDS) (**Figure 1A, Table 1**).

**Table 1:** Clinical and laboratory characteristics of the healthy controls and the COVID-19 patients.

|  | Healthy | Mild cases | Severe cases |
| --- | --- | --- | --- |
| Number at sampling | 22 | 28 | 38 |
| Median age (median (IQR<sup>a</sup>) [yrs]) | 32.50 (29.25-48) | 50.5 (34.50-60.25) | 67.5 (59.0-79.0)** |
| Gender (m/f) | 11/11 | 12/16 | 24/14 |
| Time since symptom onset (days) | - | 12.86 ± 10.71 | 20.21 ± 11.96* |
| **COVID-19 disease severity sampling / max severity<sup>b</sup>** |  |  |  |
| Mild illness– no. | - | 18 / 16 | - |
| Mild pneumonia – no. | - | 10 / 8 | - |
| Severe pneumonia – no. | - | - | 20 / 19 |
| Mild ARDS – no. | - | - | 7 / 7 |
| Moderate ARDS – no. | - | - | 7 / 8 |
| Severe ARDS – no. | - | - | 4 / 8 |
| **Laboratory values** |  |  |  |
| C-reactive protein (mean ± SD) | 1.21 ± 1.61 | 29.87 ± 51.60* | 89.46 ± 81.23** |
| Lactate dehydrogenase (% > upper limit of normal) | 0% | 28% | 73.53 %* |
| Hemoglobin (mean ± SD, [g/l]) | 139.88 ± 13.31 | 132.36 ± 28.33 | 128.18 ± 24.62 |
| Absolute platelet count (mean ± SD, [G/l]) | 257.25 ± 60.50 | 203 ± 67.97 | 219.74 ± 117.14 |
| Total white blood cell count (mean ± SD, [G/l]) | 5.74 ± 1.52 | 5.83 ± 2.90 | 6.65 ± 3.49 |
| Monocytes (mean ± SD, [G/l]) | 0.42 ± 0.15 | 0.50 ± 0.36 | 0.45 ± 0.34 |
| Neutrophils (mean ± SD, [G/l]) | 3.17 ± 1.03 | 3.74 ± 2.70 | 5.28 ± 3.36** |
| Eosinophils (mean ± SD, [G/l]) | 0.13 ± 0.08 | 0.04 ± 0.05* | 0.03 ± 0.07** |
| Basophils (mean ± SD, [G/l]) | 0.04 ± 0.02 | 0.02 ± 0.03* | 0.01 ± 0.02* |
| Lymphocytes (mean ± SD, [G/l]) | 1.95 ± 0.74 | 1.50 ± 0.69* | 0.81 ± 0.44** |
| CD3- CD56<sup>bright</sup> CD16<sup>dim</sup> NK cells (mean ± SD, [cells/ul]) | 11.14 ± 5.65 | 8.19 ± 4.98 | 6 ± 4.39* |
| CD3- CD56<sup>dim</sup> CD16<sup>bright</sup> NK cells (mean ± SD, [cells/ul]) | 206.29 ± 107.13 | 165.96 ± 147.96 | 162.16 ± 103.69 |
| **Level of care at blood sampling timepoint** |  |  |  |
| Outpatient – no. (%) | - | 14 (50) | - |
| Hospitalized – no. (%) | - | 14 (50) | 38 (100) |
| **Comorbidities** |  |  |  |
| Hypertension – no. (%) | - | 7 (25) | 22 (57.9)* |
| Diabetes – no. (%) | 1 (4.5) | 4 (14.3) | 12 (31.6) |
| Heart disease – no. (%) | - | 3 (10.7) | 17 (44.7)* |
| Cerebrovascular disease – no. (%) | - | 1 (3.6) | 4 (10.5) |
| Lung disease – no. (%) | - | 3 (10.7) | 6 (15.8) |
| Kidney disease – no. (%) | - | 7 (25) | 10 (26.3) |
| Malignancy – no. (%) | - | - | 4 (10.5) |
| Systemic Immunosuppression – no. (%) | - | 3 (10.7) | 4 (10.5) |
| * Indicates significance (p-value threshold <0.05) compared to the healthy, ** in the severe indicates significance in comparison to the healthy and the mild. |  |  |  |
| <sup>a</sup> IQR denotes the interquartile range |  |  |  |
| <sup>b</sup> COVID-19 severity at sampling according to WHO guidelines (World Health Organization, 2020) |  |  |  |

**Figure 1:**
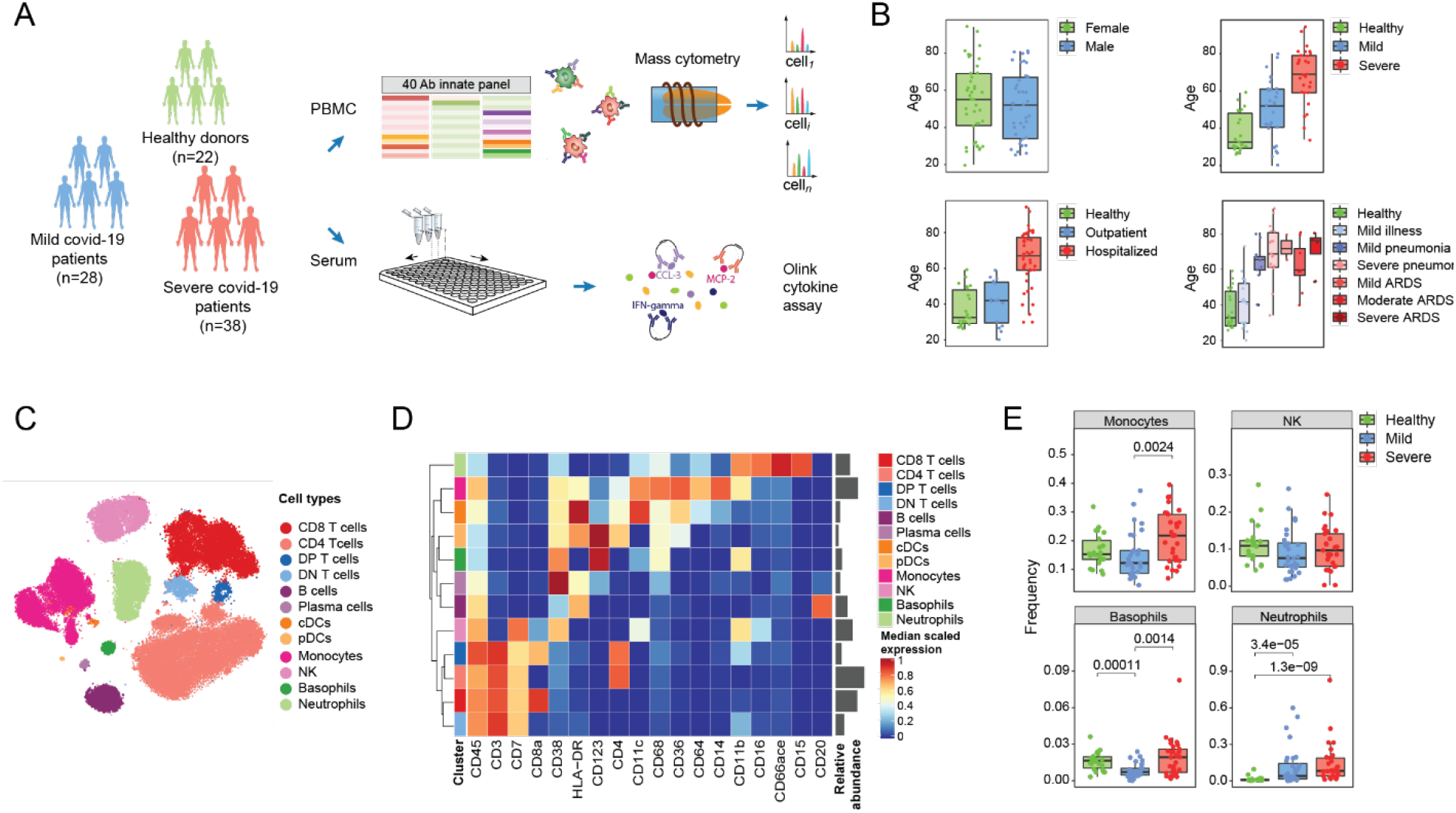
Experimental approach and identification of the main immune cell types in COVID-19 patients based on mass cytometry. (A) Schematic of the study design of a cohort of total 22 healthy controls, 28 patients with mild symptoms, and 38 COVID-19 patients with severe disease course. PBMCs of 22 healthy controls, 27 mild and 29 severe COVID-19 patients were isolated and analyzed by high-dimensional phenotypic and functional single-cell analysis by mass cytometry, and serum of 17 healthy controls, 26 mild and 36 severe COVID-19 patients was analyzed using a high-throughput multiplexed proteomics assay (Olink) detecting 92 inflammation related serum proteins. (B) Boxplots showing the age distribution in the patient cohort relative to gender, disease severity, care at sampling, and disease grade (Pneu, pneumonia; Sev Pneu, severe pneumonia; Mod ARDS, moderate ARDS; Sev ARDS, severe ARDS). (C) t-SNE plot of a random subset of 1000 immune cells from each sample colored by main cell types. (D) Heatmap of the normalized marker expression in the main cell types. Clustering was based on Euclidean distance and the Ward.D2 aggregation method. Relative abundances of each cell type are plotted to the right of the heatmap. (E) Boxplots comparing the frequencies of the indicated myeloid subsets in healthy controls and patients with mild and severe disease. p-values were calculated with a Mann-Whitney-Wilcoxon test corrected for multiple testing and are shown if the results were significant p<0.05.

We also evaluated samples from 22 healthy controls. Patients suffering from severe disease were on average older than those with mild disease (median age mild 50.5 (IQR 34.5-60.25) years, severe 67.5 (IQR 59.0-79.0) years p = 0.00032), consistent with previously published results (Wu and McGoogan, 2020) (**Figure 1B, Table 1**). Furthermore, hypertension (p = 0.0118) and heart disease (p = 0.0032) were significantly associated with a severe disease course.

The laboratory findings at admission revealed a prominent inflammatory state for patients with both mild and severe disease, as evidenced by high levels of C-reactive protein (healthy controls vs. mild p < 0.0001, mild vs. severe p < 0.0001) and pathological values of lactate dehydrogenase in mild (28%) and severe COVID-19 patients (75.53%) (p = 0.0007). The complete differential blood count showed increased neutrophil counts in patients with severe disease compared to healthy controls (5.28 ± 3.36 G/l vs. 3.17 ± 1.03 G/l, respectively (p = 0.039)). Moreover, eosinophil (healthy controls vs. mild p = 0.0004, healthy controls vs. severe p < 0.0001) and basophil counts (healthy controls vs. mild p < 0.0001, healthy controls vs. severe p < 0.0001) were significantly reduced in COVID-19 patients compared to healthy controls. Natural killer cell (NK) cytopenia in the CD3^−^ CD56^bright^CD16^dim^ population was also associated with severe disease course (healthy controls vs. severe p = 0.0023), confirming previous publications (Kuri-Cervantes *et al.*, 2020; Rodriguez *et al.*, 2020; Zhang *et al.*, 2020). The strong inflammatory state, the changes in the differential blood count and the reported prolonged clinical course before deterioration make COVID-19 a distinct disease (Giamarellos-Bourboulis *et al.*, 2020).

### Systems-wide profiling of innate compartment of patients with COVID-19

To comprehensively characterize the innate immune response against SARS-CoV-2 we took a systems-level approach (**Figure 1A**). We used a 40-plex mass cytometry (CyTOF) panel designed to identify the main immune cell types, including T cells, B cells, plasma cells, NK cells, monocytes, basophils, and neutrophils, which further allowed an in-depth characterization of myeloid cell markers and subsets (**Table S1**). In parallel, the serum levels of 92 inflammation-related proteins were quantified by targeted proteomics (**Figure 1A**). The mass cytometry dataset was acquired in two batches, using a 60-well barcoding scheme and a frozen antibody panel to minimize batch effects (**Figure S1A-D**).

The main cell types were identified using a random forest classifier trained on manually gated cells from a representative subset of data (**Figure S1E**). The cell annotation was consistent with the t-SNE map visualization (**Figure 1C, Figure S1F**) and the expression of canonical markers (**Figure 1D**). We observed that the frequencies of monocytes and basophils were increased in severe cases in comparison to mild cases and healthy controls (**Figure 1E**). Since peripheral blood mononuclear cells (PBMCs) were isolated following a density gradient separation, only low-density neutrophils were included in the analysis. Consistent with a previous report (Morrissey *et al.*, 2020) this subset was present at very low frequency in healthy controls but increased in patients infected with SARS-CoV-2, accounting for more than 50% of the PBMCs in some patients (**Figure 1E**).

### Different myeloid landscape in patients with mild and severe COVID-19

Neutrophils have been previously reported to play a key role in the development of severe forms of SARS-CoV-2 infection. In particular, a high neutrophil-to-lymphocyte ratio is associated with poor clinical outcomes, and a CD16^int^CD44^low^CD11b^int^ low-density neutrophil population, associated with high IL-6 and TNF levels, was increased in severe COVID-19 patients compared to healthy controls (Morrissey *et al.*, 2020). To assess the low-density neutrophil subsets in our cohort, we used t-SNE to visualize the expression of relevant markers on this cell type (**Figure 2A**). Although all cells were positive for the canonical neutrophil markers CD15 and CD66ace, differences in abundance of CD11b, CD11c, and CD16 were observed. Visualizing the disease status on the t-SNE map revealed an enrichment of CD16^low^ neutrophils in patients with severe disease (**Figure 2B**). To confirm this observation, we classified neutrophils into CD16^hi^, CD16^int^, and CD16^low^ populations based on manual annotation of PhenoGraph clusters (**Figure S2A-B**). The proportion of CD16^low^ neutrophils was much higher in patients with severe disease than in the other two groups (**Figure 2C**), also consistent with previous observations (Morrissey *et al.*, 2020). The CD16^low^ subset was associated with an increased proliferation rate, based on Ki-67 positivity (**Figure 2A**). Since mature neutrophils do not proliferate in the periphery, this observation indicates the release of immature or pre-mature neutrophils into the circulation, consistent with their CD11b^low^CD16^low^ phenotype (Scapini *et al.*, 2016; Ng, Ostuni and Hidalgo, 2019).

**Figure 2:**
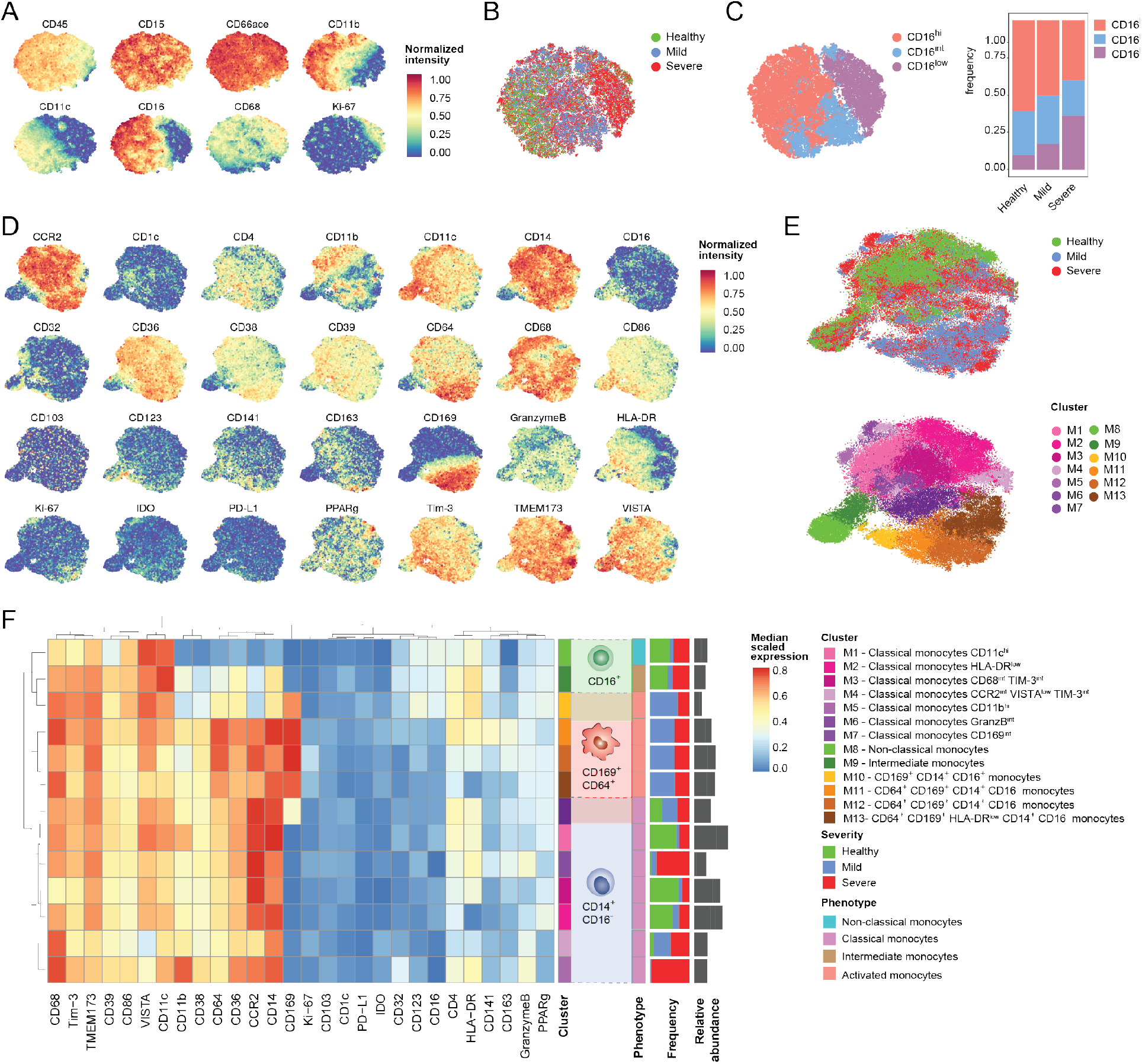
In-depth characterization of the myeloid cells in the peripheral blood of COVID-19 patients. (A) t-SNE plots of normalized expression of the indicated markers across a maximum of 1000 neutrophils per patient. (B) t-SNE plot of normalized expression of the indicated markers across a maximum of 1000 neutrophils per patient colored by disease severity. (C) Left: t-SNE plot colored by CD16 expression level based on manual assignment of PhenoGraph clusters. Right: Histogram of the proportions of CD16^hi^, CD16^Int^, and CD16^low^ neutrophils in COVID-19 patients and healthy controls. (D) t-SNE plots of normalized expression of the indicated markers across a maximum of 1000 monocytes per patient. (E) t-SNE plots as in (D) colored by disease severity (top) and by clusters identified with the PhenoGraph algorithm (bottom). (F) Heatmap of the normalized marker expression in PhenoGraph monocyte clusters. The frequency of each cluster in patients with mild and severe disease and in healthy controls is indicated. Cell numbers for each cluster are plotted to the right of the heatmap. Clusters were manually annotated to indicate phenotypes (classical, non-classical, intermediate, and activated) based on marker expression.

To characterize the phenotypic diversity in the monocytic compartment across the cohort, we visualized all myeloid-related markers included in our panel on the corresponding t-SNE map of the monocyte population (**Figure 2D**). Based on this approach, CD169, CCR2, CD14, and CD16 distinguished different monocyte subsets. We further performed automated clustering using the PhenoGraph algorithm to identify monocyte subsets in an unsupervised way, identifying 13 distinct cell communities (**Figure 2E**). Clusters M1-7 were characterized by high abundance of CD14 and CCR2 and absence of CD16, corresponding to classical monocytes, clusters M8 and M9 were CD16^+^ and M10 to M13 all showed an activated CD169^+^ phenotype.

Clusters M1-3 were found predominantly in healthy controls and were characterized by relatively low abundance of HLA-DR, CD68, and granzyme B, indicating a non-activated phenotype (**Figure 2E-F**). Clusters M4-6 were found mainly in SARS-CoV-2-infected patients and had higher levels of granzyme B compared to other classical monocytes; granzyme B production has been suggested upon TLR8 activation and enhancing FcγR-mediated antibody dependent cellular cytotoxicity (ADCC) in monocytes (Elavazhagan *et al.*, 2015). M4 additionally showed decreased levels of CCR2, TIM-3 and VISTA, which has been implicated in chemotactic paralysis in a murine model (Broughton *et al.*, 2019). Cluster M7, which expressed intermediate levels of CD169, could constitute a stage between classical monocytes and CD169^+^ classical monocytes. Cluster M9, which represented intermediate monocytes (CD14^+^CD16^+^), were found both in healthy subjects and patients with severe COVID-19, as were cluster M8 non-classical monocytes (CD14^dim^ CD16^+^).

Strikingly, CD169^+^ activated monocytes (clusters M10-13) were found exclusively in SARS-CoV-2-infected patients (**Figure 2D-E**). Cells in cluster M10 expressed CD16 and had reduced abundance of CD14 and CCR2 compared to M11-M13; these cells could be transitioning between non-classical/intermediate monocytes and the CD169^+^ compartment (**Figure 2F**). This hypothesis is further supported by a diffusion map analysis (Haghverdi, Buettner and Theis, 2015), which aligns cells along putative developmental trajectories and suggests that CD169^+^ could derive from both the CD16^+^ and the CD16^−^ compartment (**Figure S2C, D**). Most markers, including CD64, CD169, CD4, and HLA-DR were progressively decreasing in clusters M11 to M13, suggesting that these clusters were part of a phenotypic continuum rather than representing distinct cell subsets. These clusters were also characterized by increased Ki-67 positivity, especially M12, compared to non-classical, intermediate or classical monocytes (**Figure 2F**). In summary, high-dimensional single-cell mass cytometry analysis allowed us to characterize the monocyte compartments of COVID-19 patients and healthy controls with unprecedented depth and to uncover profound changes upon SARS-CoV-2 infection.

### Stratification of COVID-19 patients based on monocyte composition

We next assessed the distribution of the 13 identified monocyte clusters across patients. We calculated the frequencies of each of the 13 clusters on a per sample basis and performed hierarchical clustering to order the patients by compositional similarities (**Figure 3A**). This analysis revealed three main groups, which were enriched for mild cases (**Figure 3A**, left), healthy controls (middle), and severe cases of COVID-19 (right). In the group that included mostly mild cases, the monocyte compartment consisted almost exclusively of the CD169^+^ clusters (M10-13) in different ratios, with only a minor fraction of cells from clusters M4-7. The healthy controls were relatively homogenous: About 80% of cells consisted of classical monocytes (clusters M1-3) and about 15-20% of cells were intermediate and non-classical monocytes (clusters M9 and M8, respectively) consistent with the literature (Thomas *et al.*, 2017). The group dominated by patients with severe COVID-19 was characterized by a high frequency of distinct classical monocyte subsets (clusters M4-7), with frequencies of intermediate and non-classical monocytes slightly higher than in the group dominated by healthy controls.

**Figure 3:**
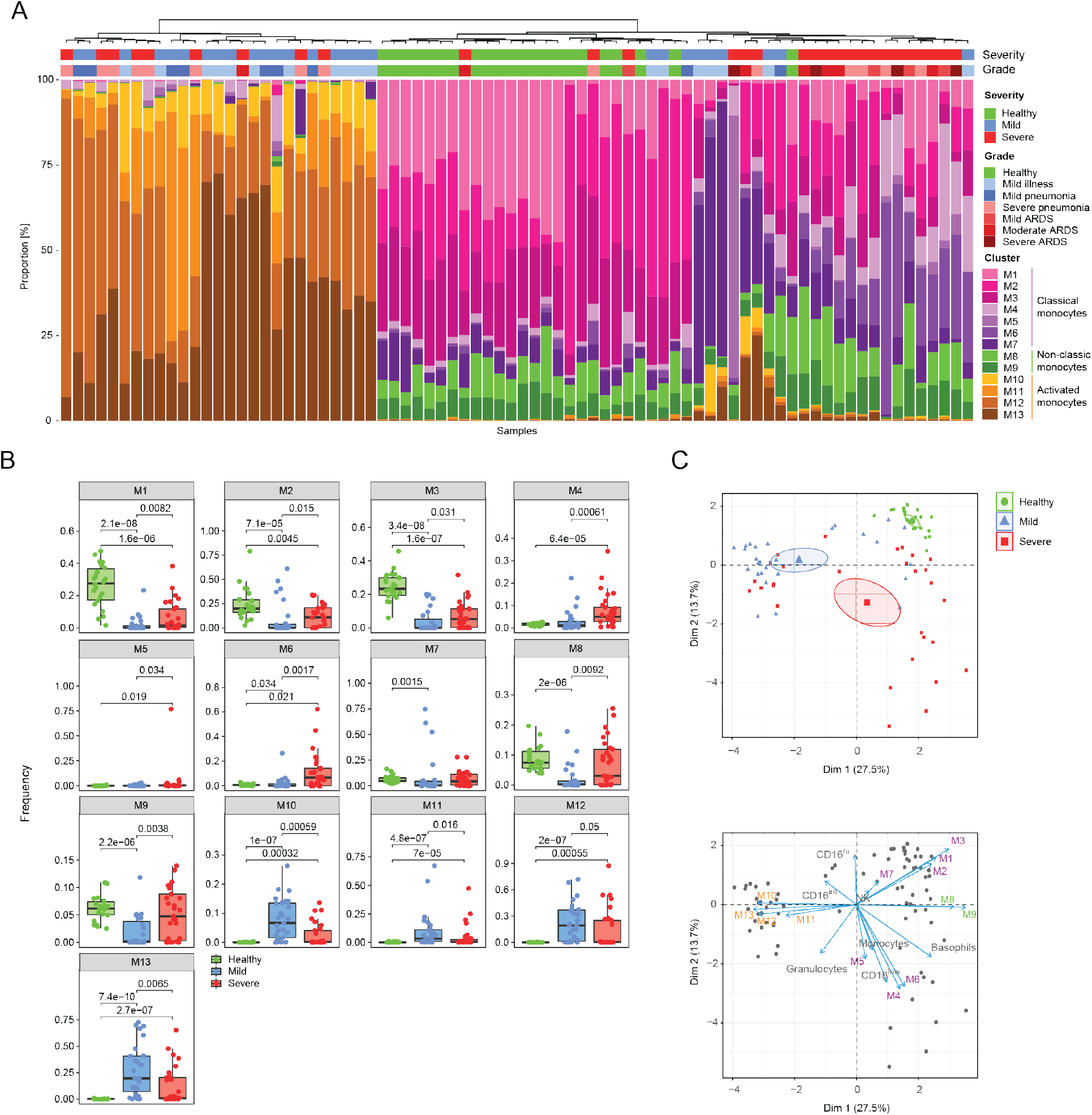
Patient stratification based on myeloid signature. (A) Stacked histogram of the PhenoGraph monocyte clusters per patient, ordered by cluster composition similarities based on Euclidean distance. Disease severity and grade for each patient are shown. (B) Boxplots of frequencies of the indicated monocyte clusters in the different disease severities. p-values were calculated with a Mann -Whitney-Wilcoxon test corrected for multiple testing and are shown if the results were significant p<0.05. (C) Top: Principal component analysis (PCA) of monocyte and neutrophil cluster frequencies and myeloid immune cell subset frequencies across the cohort. The PCA plot (top) shows the two first principal components (PCs) separating the samples. Each dot represents a patient, colored by disease status. Bottom: Biplot of parameters contributing to the separation. Arrow lengths and directions in the biplot indicate the importance of the parameter to the PCs.

We observed similar patterns when we directly compared frequency differences of the different monocyte clusters in healthy subjects and patients with mild and severe disease (**Figure 3B**). Most strikingly, the CD169^+^ clusters M10-13 were completely absent in healthy controls and were significantly higher in patients with mild disease than those with severe disease, with the exception of cluster M12 where the significance level was not reached. Conversely, the classical monocyte clusters M1-3 were present at high levels in healthy donors, at lower levels in the patients with mild disease, and at intermediate levels in patients with severe disease. Clusters M4-6, also defined as classical monocytes, were virtually absent in healthy controls and were present at higher levels in patients with severe than mild disease. The non-classical (M8) and the intermediate monocytes (M9) were significantly reduced in patients with mild disease compared to healthy controls and were present at higher levels in a subset of patients with severe disease than in healthy controls.

To gain more insight into the relationship between the innate immune signature and disease status, we performed a principal component analysis (PCA) of monocyte and neutrophil cluster frequencies and myeloid immune cell subset frequencies across the cohort and asked whether these innate immune signatures varied with disease severity. Indeed, the first two principal components enabled the stratification of subjects based on disease status (**Figure 3C, left panel**). A biplot graph displaying simultaneously the subjects and the eigenvectors of the different cell subsets revealed a strong association between clusters M1-M3 and healthy controls (**Figure 3C, right panel**). A group dominated by mild patients was characterized by high levels of M10 to M13 clusters. A more heterogeneous set of predominantly severe COVID-19 cases were defined by higher levels of M4-M6 and M8-M9 clusters and by CD16^low^ low-density neutrophils. A correlative analysis performed across innate cell subsets and patients confirmed the pattern observed based on the PCA analysis (**Figure S3A**). Thus, despite the expected diversity across individuals, these multiparametric analysis identified innate immune signatures that allowed a stratification of healthy donors, patients with mild COVID-19, and patients with severe disease.

### Changes in innate cell frequencies over the course of SARS-CoV-2 infection

We used the fact that the patients presented to hospitals at different times after symptom onset to examine cell cluster frequencies over the disease course. This allowed us to gain an understanding of the temporal dynamics of the different immune subsets present during SARS-CoV-2 infection. The total monocyte compartment remained relatively constant over the disease course, but monocytes were present at higher frequencies in patients with severe compared to mild disease (**Figure 4A**). The low-density neutrophils were present at higher frequency in COVID-19 patients compared to healthy controls early after symptom onset and decreased at later stages of the disease. These changes were accompanied by a decrease of CD16^hi^ neutrophils over disease course in patient with severe disease, whereas CD16^low^ neutrophils remained consistently high (**Figure 4B**).

**Figure 4:**
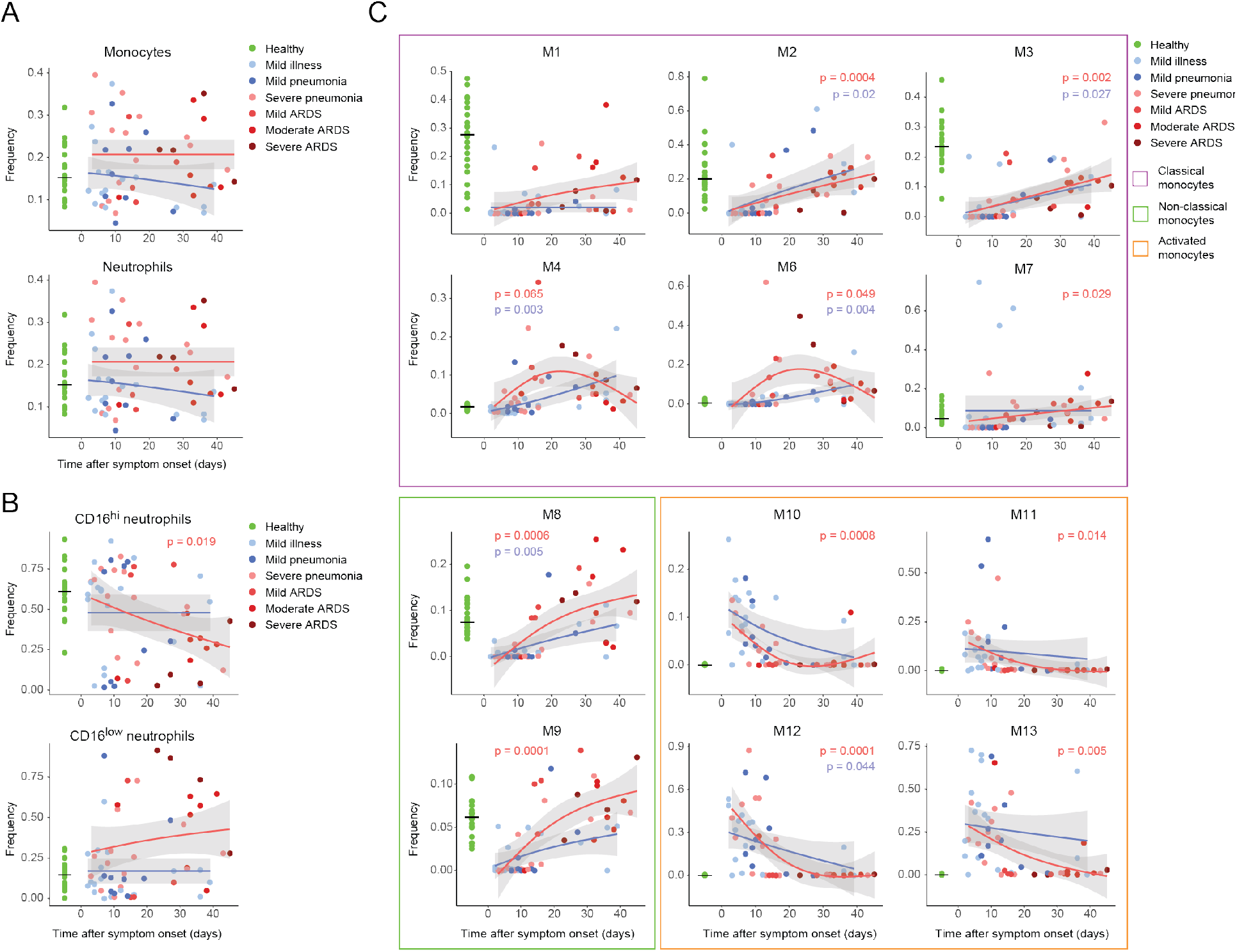
Myeloid cell frequencies over the course of the SARS-CoV-2 infection. (A) Scatter plot of monocyte and granulocyte frequencies relative to the time after symptom onset. The dots are colored by disease grade at sampling time. The frequencies in healthy controls are shown as a reference on the left. The pseudo-time course was modeled using a general additive model for the disease severities separately (mild, blue lines; severe, red lines). (B) Scatter plot of indicated neutrophil subset frequencies relative to the time after symptom onset. The cluster frequency is given in relation to the total neutrophils in the PBMCs. (C) Scatter plot of indicated monocyte subset frequencies relative to the time after symptom onset. The cluster frequency is given in relation to the total monocytes in the PBMCs. The clusters associated with non-classical, classical, and activated monocytes are indicated by purple, green, and orange boxes, respectively.

The monocyte subsets (clusters M1-13) also showed distinct changes over the disease course. Compared to healthy controls, clusters M1, M2 and M3 were observed at very low frequencies in mild and severe cases sampled early during the disease course but gradually increased in patients sampled later during their disease course (**Figure 4C, purple box**). The frequencies of these clusters tended to remain at lower levels than in the healthy controls even at time points as late as 47 days after symptom onset. (**Figure 4C, purple box**). In contrast, frequencies of clusters M4 and M6, which were not present in the healthy controls, increased during the disease course, more notably in severe cases of COVID-19, with a peak around three weeks after symptom onset. (**Figure 4C, purple box**). Intriguingly, mild COVID-19 cases with increased frequencies of cluster M4 were mostly hospitalized, further progressed to severe COVID-19 at a later time point or showed prolonged symptoms for more than two months (**Figure S4A**).

The activated CD169^+^ monocyte clusters M10-M13 were present at high frequencies early during the disease course (**Figure 4C, orange box**). Within the first 20 days after symptom onset, the monocyte compartment was strongly dominated by this phenotype, which decreased thereafter and was undetectable in most patients sampled more than 20 days after symptom onset. This pattern was similar in both groups with mild and severe COVID-19.

The CD16^+^CD14^dim^ M8 (non-classical) and CD16^+^CD14^+^ M9 (intermediate) monocyte clusters were virtually absent at early stage upon viral infection (**Figure 4C, green box**), consistent with data from single cell-RNA sequencing studies on PBMCs (Wilk *et al.*, 2020). The initial reduction was followed by their recovery at later stages of disease, showing a trend towards higher frequencies in patients with severe compared to mild disease for both M8 and M9.

In summary, the analysis of cluster frequencies in COVID-19 patients in relation to the time since symptom onset revealed distinct dynamic patterns of monocyte frequencies in patients with mild and severe disease. At early stages in all patients, the monocyte compartment was dominated by activated monocytes (cluster M10-13), which was characteristic for COVID-19 patients. Despite a normalization of the CD14^+^ monocyte compartment in the blood thereafter, the composition of the monocyte compartment differed from that of healthy subjects 40 days after symptom onset in both groups with mild and severe COVID-19.

### Temporal correlation of cytokine signatures and innate cell subsets

Monocyte development, homeostasis, and fate are strongly interlinked with the cytokine and chemokine environment. To probe potential correlates to the changes seen in the myeloid compartment during COVID-19 progression, we used the targeted proteomics Olink assay to measure 92 inflammation-associated serum proteins in samples from 17 healthy controls, 26 patients with mild disease, and 36 patients with severe disease who were concomitantly analyzed by mass cytometry. The comparison of data from healthy controls and patients with severe COVID-19 showed a strong upregulation of proinflammatory cytokines and chemokines (**Figure 5A**). IL-6, TNF, IFNγ, and IL-18 were significantly upregulated (significance cut-off false discovery rate (FDR) 1%) in patients with severe disease as shown by the proteomics measurements (**Figure 5A)** consistent with previous reports (Blanco-Melo *et al.*, 2020; Veerdonk *et al.*, 2020). The elevated levels of IL-18 and increased LDH we observed might indicate a contribution of inflammasome activation (Lee *et al.*, 2020; Merad and Martin, 2020; Yap, Moriyama and Iwasaki, 2020), although we did not detect an elevation of IL-1β (**Figure S5A**).

**Figure 5:**
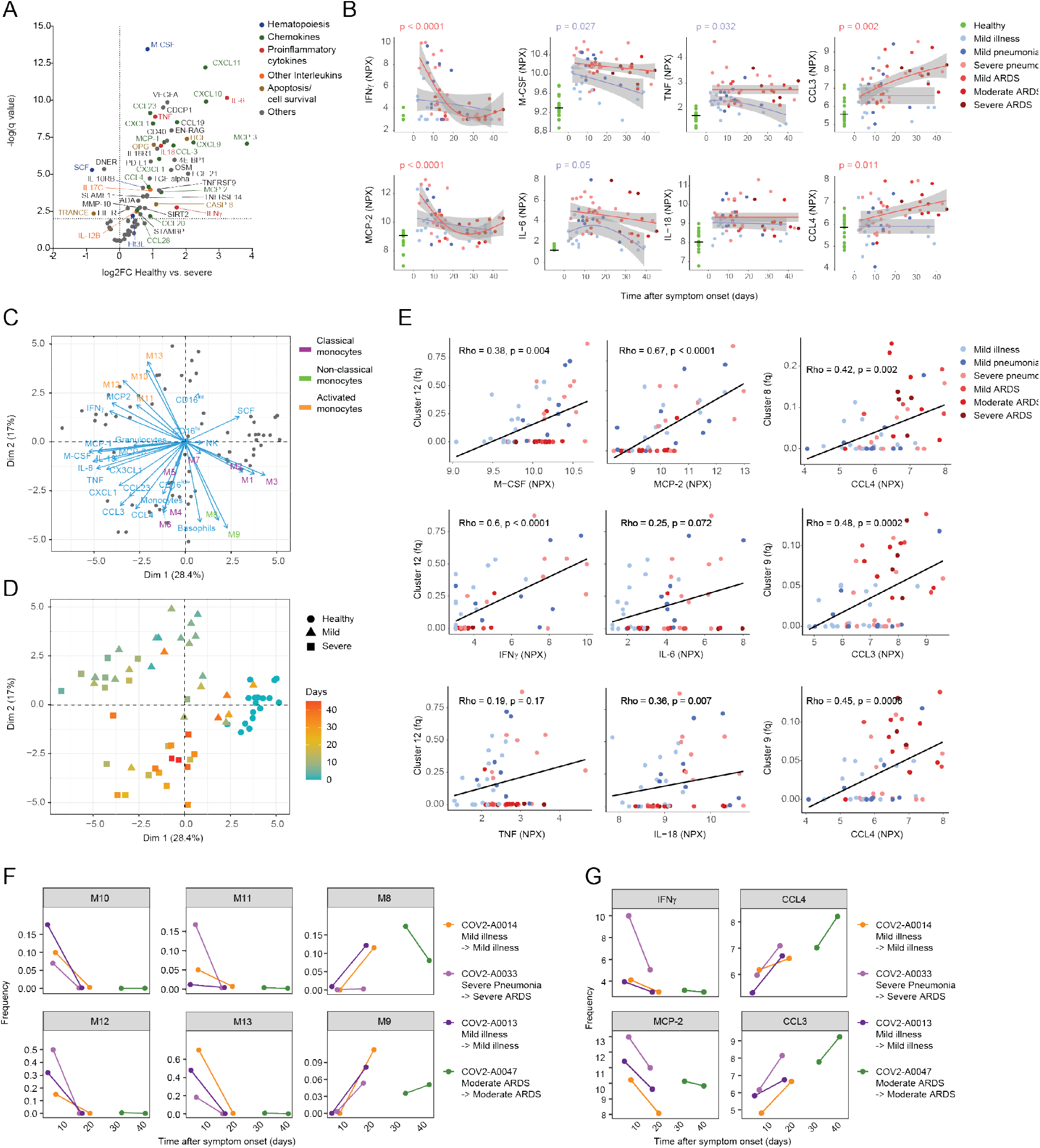
Cytokine signature shift between early and late stages of disease correlates with innate cell subsets. (A) Volcano plot of the Olink proteomics data comparing the healthy control data to that from patients with severe COVID-19. An FDR of 1% was taken as significance cut-off. (B) Scatter plot of serum protein expression levels relative to the time after symptom onset. Plotted is normalized protein expression (NPX) on a log2 scale. The dots are colored by disease grade at sampling time. The expression levels of the healthy controls are shown as a reference on the left. The pseudo-time course was modeled using a general additive model for the disease severities separately (mild, blue lines; severe, red lines). *Legend continues on next page* (C) Biplot of the first two PCs of monocyte and neutrophil cluster frequencies, myeloid immune cell subset frequencies, and expression values of selected serum proteins. Dots represent the COVID-19 patients and healthy controls, and the arrow lengths and directions indicate the importance of the parameter to the PCs. (D) Scatter plot of the first two PCs of monocyte and neutrophil cluster frequencies, and myeloid immune cell subset frequencies, and expression values of selected serum proteins colored by the time since symptom onset. (E) Scatter plots of frequencies of the indicated clusters versus expression of selected serum proteins in individual patients. The dots indicate data for individual patients colored by disease grade. Relationship between the two variables is visualized with a linear regression line and quantified using a Spearman’s correlation coefficient (Rho) with the corresponding p value. (F) Scatter plots of frequencies of the indicated monocyte clusters in individual patients who were sampled twice during the course of the study. Dots are colored by patient number, and lines connect paired samples. (G) Scatter plots of NPX on a log2 scale of the indicated soluble factors in patients who were sampled twice during the course of the study. Dots are colored by patient number, and lines connect paired samples.

Several chemokines important for myeloid cell trafficking were significantly upregulated in patients with severe COVID-19. These chemokines included MCP-1 (also known as CCL2), MCP-2 (also known as CCL8), MCP-3 (also known as CCL7), CX3CL1 (chemotactic for non-classical monocytes), CXCL1 (a potent neutrophil-recruiting chemokine), and the more promiscuous CCL3 and CCL4. M-CSF, which is crucial for myeloid precursor survival and lineage commitment (Mossadegh-Keller *et al.*, 2013; Boettcher and Manz, 2017), was significantly increased in severe as well as mild COVID-19 patients compared to healthy controls (**Figure 5A, Figure S5B**).

Distinct expression patterns emerged by plotting cytokine expression versus time after symptom onset. The relation was modeled using a general additive model as in Figure 4. In comparison to healthy controls, IFNγ and MCP-2 were present at significantly higher concentrations in patients with severe disease sampled early after symptom onset but returned to near-normal values at later stages of the disease course. Consistent with confirmatory ELISA measurements, M-CSF, IL-6 and TNF, which were all present at higher levels in patients with severe disease than mild disease, tended to be present at high levels throughout the disease course in patients with severe disease, whereas a decrease was observed in patients with mild symptoms as the disease progressed (**Figure 5B, Figure S5A**). The chemokines CXCL1, CX3CL1, MCP-1, and MCP-3 had significantly increased levels compared to healthy controls in patients with severe COVID-19 independent of time after symptom onset (**Figure S5B**). In striking contrast, CCL3 and CCL4 were not induced in patients in the first days of the disease but were significantly increased at later stages in patients with severe disease (**Figure 5B)**. This is suggestive of a more exacerbated inflammatory phenotype late during the disease course in patients with severe disease than in those with mild symptoms. Differential expression analysis confirmed this observation (**Figure S5C)**.

To better understand the interplay between these serum proteins and cell subsets of the myeloid compartment, we performed a hierarchical clustering on a correlation map of all significantly changed (FDR 1%) serum proteins and immune cell subsets and myeloid clusters (**Figure S5D**). This approach demonstrated a strong association between the activated CD169^+^ monocyte clusters M10-13 with pro-inflammatory cytokines, including IFNγ, MCP-2, IL-18, and IL-6, and an association between the classical monocyte clusters M4 and M5 with TGF-α, CCL3, and CCL4. To further understand these relationships in the context of the patients in our cohort, we performed PCA. This analysis revealed that certain combinations of cellular and soluble factors stratified patients and healthy controls (**Figure 5C, D**). One group, dominated by patients with mild symptoms and patients sampled early in the disease course, was defined by the clusters M10-13 in combination with the cytokines MCP-2 and IFNγ. A second group of patients, enriched for more severe, late-stage cases, was defined primarily by the non-classical clusters M4-6 in combination with the cytokines CCL3, CCL4, and CCL23 (**Figure 5C, D**). The healthy donors were defined by the clusters M1-3 and were located close to the majority of late-stage mild patients. The intermediate and non-classical clusters M8 and M9 contributed to both the healthy and the severe groups. These data strongly suggest an innate signature shift between the early and the late stage of the disease, leading to a divergence of patients with mild and severe COVID-19 over the disease course. While the former became similar to healthy controls, the latter exhibited signs of hyper-inflammation.

To further characterize the relationships between cell clusters and soluble factors identified in the PCA, we performed a direct correlation between the cluster frequencies and cytokine levels. There was a strong correlation between cluster M12 and M-CSF, IFNγ, IL-6, and TNF, which was most evident in early-stage patients (**Figure 5E, Figure S5E)**. We also found that the non-classical monocytes M8 and the intermediate monocyte cluster M9 were strongly correlated with CCL3 and CCL4, but in this case the correlation was most prominent in late-stage patients (**Figure 5E, right panels, Figure S5F**). These results suggest that the inflammatory environment found in late stages of COVID-19 is predominantly associated with the presence of the intermediate monocytes.

To determine whether the signature switch observed in our pseudo-temporal analysis was observed in individual patients over time, we compared the cluster frequencies and cytokine levels in four patients who were sampled twice. We confirmed that the frequencies of the activated CD169^+^ clusters M10-13 were reduced over time, and the frequencies of the non-classical monocyte cluster M8 and the intermediate monocyte cluster M9 were increased (**Figure 5F**). Similarly, we again observed decreases of IFNγ and MCP-2 over time in these re-sampled patients, accompanied by increases in CCL3 and CCL4, providing strong support to the findings made based on the pseudo-temporal analysis (**Figure 5G**). Overall, these data suggest that there is a coordinated change from a CD169+, IFNγ, and MCP-2 phenotype to an intermediate monocyte, CCL3, and CCL4 phenotype in the first two weeks after SARS-CoV-2 infection and that patients with severe COVID-19, who show long-lasting high levels of CCL3 and CCL4, show a more dramatic phenotypic switch.

## Discussion

Early in the COVID-19 pandemic, data began to suggest that patients with severe disease have hyperinflammatory immune responses with changes in the myeloid compartment toward a pro-inflammatory phenotype (Huang *et al.*, 2020; Liao *et al.*, 2020; Wilk *et al.*, 2020). Employing a systems-wide immune characterization on a multicenter cohort of COVID-19 patients and healthy controls we identified marked changes in the innate immune signature in SARS-CoV-2 infected individuals.

We discovered that the myeloid compartment undergoes profound phenotype changes during a SARS-CoV-2 infection with a decrease in the classical monocyte clusters M1, M2 and M3, a depletion of the intermediate and non-classical monocyte clusters M9 and M8, toward a CD169^+^ activated monocyte phenotype and a surge of low-density neutrophils early in the disease course. The immature phenotype of the neutrophils is in accordance with data in whole blood (Carissimo *et al.*, 2020).

CD169 is a sialoadhesin involved in pathogen uptake, which is quickly induced in a type I IFN dependent manner on the surface of monocytes upon Epstein–Barr virus (EBV) or human immunodeficiency virus (HIV) infection (Rempel *et al.*, 2008; Farina *et al.*, 2017). Consistent with our findings, CD169 has been reported on circulating monocytes in COVID-19 patients (Bedin *et al.*, 2020; Carissimo *et al.*, 2020), while type I IFNs have been shown to be a hallmark of SARS-CoV-2 infection and an impaired type I IFN response has been linked with severe disease (Hadjadj *et al.*, 2020; Wilk *et al.*, 2020).

Our data suggest that CD169^+^ monocytes arise from classical, intermediate and non-classical monocytes, similarly to what was reported upon simian immunodeficiency virus (SIV) or HIV infection (van der Kuyl *et al.*, 2007; Kim *et al.*, 2015). The fate of these CD169^+^ monocytes, which disappear quickly from the blood, remain elusive. Bronchioalveolar lavage fluid and SARS-CoV mouse models have shown that inflammatory monocyte-derived macrophages play crucial roles in local inflammation and tissue damage (Channappanavar *et al.*, 2016; Liao *et al.*, 2020). Recruitment of these activated CD169^+^CCR2^+^ monocytes to the lung through the CCR2-CCL2 axis is possible as MCP-1 (CCL2) is significantly upregulated in COVID-19 patients. These cells could then contribute to the lung damage observed in patients with severe disease.

Analyzing the inflammatory cytokine and chemokine response to SARS-CoV-2 we identified a TNF and IL-6-dominated inflammatory response in COVID-19 patients in agreement with a previous report (Veerdonk *et al.*, 2020). The initial inflammatory response was dominated by IFNγ, MCP-2, M-CSF, and IL-6, which were coregulated with the CD169^+^ monocyte subsets, most prominently the M12 cluster, and the low-density granulocytes as evidenced by an unsupervised correlation matrix. IL-6 and M-CSF contribute to increased hematopoiesis under inflammatory conditions (Boettcher and Manz, 2017), potentially explaining the initial increase of activated monocytes and granulocytes after SARS-CoV-2 infection. Monocytes and neutrophils are generally considered to lose their proliferative capacity once they leave the bone marrow, a process driven by the chemokines MCP-1 and MCP-3 (Swirski, Hilgendorf and Robbins, 2014). A small subset of monocytes has been shown to retain proliferative capacity especially under inflammatory conditions. These cells might contribute to the initial monocyte wave as we detected increased Ki-67 positivity in these cells. However, this might also be explained by a more immature phenotype of these cells (Clanchy, 2006; Patel *et al.*, 2017). Our proteomics panel did not include G-CSF and GM-CSF, two other cytokines crucially involved in myeloid homeostasis (Lang *et al.*, 2020).

Late in the disease course, COVID-19 patients with severe disease continued to show signs of ongoing inflammation, including abnormally high levels of TNF. The innate signature at this stage was driven by a surge of the disease specific clusters M4-M6, the reappearance of intermediate and non-classical monocytes, which reached particularly high levels in severe COVID-19 patients, and the chemokines CCL3 and CCL4. These CCR5 ligands drive the recruitment of a variety of immune cells such as neutrophils and monocytes, but also adaptive immune cells, and are frequently increased in acute respiratory viral infections (Glass, Rosenberg and Murphy, 2003; Nuriev and Johansson, 2019). Both have been shown to be produced on a transcriptomic level by monocytes in COVID-19 patients (Lee *et al.*, 2020; Liao *et al.*, 2020). It could be hypothesized that the production of these chemokines at late disease stages in severe COVID-19 is a correlate of ongoing local viral replication leading subsequently to a continuous systemic inflammatory state (Lucas *et al.*, 2020), potentially as a result of an inadequate T cell response (Adamo *et al.*, 2020).

CD16^+^ monocytes have gained considerable attention in the pathophysiology of COVID-19 as they are critical for viral sensing and the subsequent inflammatory response, including via the production of CCL3 and CCL4 (Cros *et al.*, 2010; Kwissa *et al.*, 2014; Zhou *et al.*, 2020). The strong reduction of CD16^+^ monocytes M8 and M9 early in the disease course is striking. Several mechanisms such as differentiation, ie. acquisition of an inflammatory phenotype, or migration, potentially dependent on CX3CL1 (Thomas *et al.*, 2015), could be involved. A recent study in bronchoscopy samples from COVID-19 patients in the ICU found an enrichment of CD16^+^ monocytes in the lung (Sanchez-Cerrillo *et al.*, 2020).

This raises a caveat of our study, as we analyzed only peripheral blood samples. Another limitation is that only four patients were analyzed longitudinally, whereas the cellular trajectories during the acute infection rely on samples collected from multiple individuals who presented at different times after symptom onset. However, our time course analysis also highlights the importance of the sampling time point in analyzing the immune response. Notably, the paired sample analysis confirmed the patterns observed at the cohort level, providing strong support to the pseudo-time analysis.

In summary, our systems-level analysis of the innate immune response to SARS-CoV-2 shows that there are profound changes in the peripheral monocyte compartment that are largely similar in cases of mild and severe disease. However, the patients with severe symptoms have a markedly stronger inflammatory phenotype throughout the disease course and most prominently show a distinct innate signature at later stages of the disease. These results provide evidence for a strong inflammatory response to SARS-CoV-2 infection, further supporting investigation of targeted anti-inflammatory interventions in severe cases of COVID-19 (Merad and Martin, 2020). The distinct time-dependent change in immune signatures indicate that specific interventions might benefit from precise timing to maximize therapeutic efficacy (Lang *et al.*, 2020).

## Methods

### Human subjects and patient characteristics

Patients were recruited at the Hospital Uster, Hospital Limmattal, Triemli Hospital, and the University Hospital Zurich (Switzerland) from an outpatient as well as inpatient setting. The patients were eligible if they were symptomatic at the time of inclusion, had a newly diagnosed SARS-CoV-2 infection confirmed by quantitative reverse-transcriptase polymerase chain reaction (RT-qPCR), and were more than 18 years old. Healthy donors (n=22) were recruited as controls. All participants, patients and healthy controls, signed a written informed consent. The study was approved by the Cantonal Ethics Committee of Zurich (BASEC #2016-01440) and performed in accordance with the Declaration of Helsinki.

Standard clinical laboratory data (CRP, LDH, complete blood count with differential) was collected from the first day of hospitalization until the end of hospitalization. Patients were classified according to WHO criteria (World Health Organization, 2020) into mild cases (those with mild illness and mild pneumonia) and (b) severe cases (those with severe pneumonia and ARDS, as defined by the Berlin definition (Ranieri *et al.*, 2012)). A blood sample was collected from each patient, if possible coordinated with the usual care. For longitudinal analysis of SARS-CoV-2-specific immune responses two subjects with mild COVID-19 and two subjects with severe COVID-19 were sampled twice during their disease course. All samples were processed in the same hospital laboratory.

In total 70 COVID-19 patients and 22 healthy subjects were recruited. Four patients were excluded from analysis (two due to chronic lymphocytic leukemia and two because it was unclear whether the current disease was the primary infection). All healthy controls were tested for SARS-CoV-2 specific IgA and IgG antibodies and all were below the diagnostic reference value. The complete characteristics of the cohort are given in **Table 1**.

All patients received a standard clinical laboratory sampling and cytokines were measured. Furthermore, samples from 27 COVID-19 patients with mild, 29 with severe disease and all healthy subjects were processed for CyTOF. Samples from 26 COVID-19 patients with mild, 36 with severe disease and 17 healthy patients were evaluated with Olink proteomics. 54 patients could be analyzed by CyTOF and Olink. Longitudinal samples from patient COV2-A0013 homogeneously failed the Olink incubation control, and could thus only be compared with each other, but were excluded from other analysis, together with one other sample which was not correctly processed prior to analysis. Routine flow cytometry for NK cell quantification was performed on all samples in the accredited immunological laboratory at the University Hospital Zurich, as previously described (Adamo *et al.*, 2020). The cohort characteristics and a selection of the Olink dataset are also shown in Adamo et al. describing the T cell response of this cohort.

### Blood collection and sample preparation for CyTOF

Venous blood samples were collected in BD vacutainer EDTA tubes, centrifuged, plasma removed, and the remaining blood diluted with an equal amount of PBS. This mixture was then layered into a SepMate tube (STEMCELL, Cat. #85460) filled with lymphodex (Inno-Train Diagnostik GmBH, Cat. #002041500) solution. The tube was centrifuged, and the PBMCs were washed with PBS and re-centrifuged. Aliquots of 1×10^6^ PBMCs were then centrifuged, resuspended in 200 μL 1.6% PFA (Electron Microscopy Sciences) diluted with RPMI 1640 medium, and fixed at room temperature for 10 min. Subsequently the reaction was stopped by adding 1 mL of cell staining medium (CSM, PBS with 0.5% bovine serum albumin and 0.02% sodium azide). The cells were centrifuged and the disrupted pellet was frozen at −80 °C. The remaining PBMCs were frozen and stored in 1.5 mL 90% FBS, 10% DMSO at −80 °C for at least 4 h. For long-term storage, the frozen cells were moved to the liquid nitrogen. The reference cells for the CyTOF analysis, PBMCs from a healthy donor, were stimulated with either 0.1 μg/mL phytohemagglutinin for 24 h or 1 μg/mL lipopolysaccharide and 1.5 μg/mL monensin for 48 h. One-third of the PBMCs were unstimulated. After stimulation or not, the PBMCs were fixed and frozen as described above.

### Mass cytometry barcoding

We ensured homogenous staining by barcoding 1 × 10^6^ PBMCs from each patient using a 60-well barcoding scheme consisting of unique combinations of four out of eight barcoding reagents as previously described (Zunder *et al.*, 2015). Six palladium isotopes (^102^Pd, ^104^Pd, ^105^Pd, ^106^Pd, ^108^Pd, and ^110^Pd, Fluidigm) were chelated to 1-(4-isothiocyanatobenzyl)ethylenediamine-N,N,N’,N’ tetraacetic acid (Dojino). Two indium isotopes (^113^In and ^115^In, Fluidigm) were chelated to 1,4,7,10-tetraazacy-clododecane-1,4,7-tris-acetic acid 10-maleimide ethylacetamide (Dojino) following standard procedures (Zivanovic, Jacobs and Bodenmiller, 2014). We titrated mass tag barcoding reagents to ensure equivalent staining for each reagent; final concentrations were between 50 nM and 200 nM. We used the previously described transient partial permeabilization approach to barcode the cells (Behbehani *et al.*, 2014). PBMCs from all samples were randomly loaded into wells of two 96-well plates and were analyzed in two independent experiments. Three standard samples were loaded onto each plate to enable assessment of inter-run variability. Cells were washed with 0.03% saponin in PBS (PBS-S, Sigma Aldrich) and incubated for 30 min with 200 μL of mass tag barcoding reagents diluted in PBS-S. After washing three times with CSM, samples from each plate were pooled and then split into two tubes for staining with the two antibody panels.

### Antibodies and antibody labeling

The antibodies used in this study, including provider, clone, and metal tag, are listed in **Table S1**. Antibody conjugation was performed using the MaxPAR antibody labelling kit (Fluidigm). Upon conjugation, the yield of recovered antibody was assessed on a Nanodrop (Thermo Scientific) and then supplemented each antibody with Candor Antibody Stabilizer. We performed titrations to determine optimal concentrations of all conjugated antibodies. All antibodies used in this study were managed using the cloud-based platform AirLab (Catena *et al.*, 2016).

### Sample staining and data acquisition

After barcoding, pooled cells were incubated with FcR blocking reagent (Miltenyi Biotec) for 10 min at 4 °C. Cells were stained with 400 μL of the antibody panel per 10^7^ cells for 45 min at 4 °C. Cells were washed three times in CSM, once in PBS, resuspended in 0.4 ml of 0.5 μM nucleic acid Ir-labeled intercalator (Fluidigm) and incubated overnight at 4 °C. Samples were then prepared for CyTOF acquisition by washing the cells once in CSM, once in PBS, and once in water. Cells were then diluted to 0.5 × 10^6^ cells/mL in Cell Acquisition Solution (Fluidigm) containing 10% EQ™ Four Element Calibration Beads (Fluidigm). Samples were acquired on a Helios upgraded CyTOF 2. Individual .fcs files collected from each set of samples were pre-processed using an semi-automated R pipeline based on CATALYST to perform individual file concatenation, bead based normalization, compensation, debarcoding, and batch correction as previously described (Crowell *et al.*, 2020). Spillover matrix for CyTOF compensation was assessed on all antibodies used in this study as previously suggested (Chevrier *et al.*, 2018).

### Mass cytometry data analysis

Upon pre-processing, a subset of 1,000 randomly selected cells from each sample were exported as FCS files and loaded on Cytobank. Immune cell subsets were manually gated according to the scheme described in **Figure S1D**. FCS files corresponding to each gate were exported and used to train a random forest classifier (R package randomForest), based on 500 trees and 6 variables tried at each split, leading to an OOB estimate of error rate of 0.43%. The resulting random forest model was used to assign each cell of the dataset to the predefined cell types. Based on a 40% assignment probability cutoff and a 20% delta cutoff, 98% of the cells were retained in the analysis. To visualize the high-dimensional data in two dimensions, the t-SNE algorithm was applied on data from a maximum of 1,000 randomly selected cells from each sample, with a perplexity set to 80, using the implementation of t-SNE available in CATALYST (Nowicka *et al.*, 2019). Channels which were not relevant for these cell subsets or which were affected by different background stainings across batches were excluded (CD15, CD66ace, CD3, CD45, CD8a, CD20, CXCR2, GranzymeB). Data were displayed using the ggplot2 R package or the plotting functions of CATALYST (Nowicka *et al.*, 2019).

Visualization of marker expression on t-SNE maps was performed upon data normalization between 0 and 1. The maximum intensity was defined as the 99th percentile. In order to perform hierarchical clustering, pairwise distances between samples were calculated using the Spearman correlation or euclidean distance, as indicated in the figure legend. Dendrograms were generated using Ward.2’s method. Heatmaps were generated based on the pheatmap package. Clustering analysis of the myeloid and neutrophil subsets was performed using the R implementation of PhenoGraph run on all samples simultaneously, with the parameter *k*, defining the number of nearest neighbors, set to 100 (Levine *et al.*, 2015). For the myeloid subset, clusters with less than 600 cells were excluded from the analysis.

To identify putative single-cell trajectories among monocyte clusters, we used the implementation of the diffusion map algorithm available in the R package scater using the default parameters and the same channels used to perform the t-SNE analysis. A maximum of 1,000 cells randomly selected from each cluster were included in the analysis.

The principal component analysis to identify the variations in the data described by the cluster frequencies or the combination of cluster frequencies and cytokine levels was performed based on the FactoMineR package. Data were visualized using the factoextra R package.

### Cytokine ELISA

Serum was collected in BD vacutainer tubes. The samples were processed in the accredited immunological laboratory at the University Hospital Zurich. IL-1β, IL-2, IL-6, IFNγ, TNFα were quantified using R&D Systems ELISA kits.

### Proteomics analysis using Olink

For serum proteomics, the commercially available proximity extension assay-based technology from Olink® Proteomics was used. Heat-inactivated plasma samples were sent to the Olink analysis laboratory in Davos, Switzerland for analysis using the inflammation panel. The Olink technology has been described previously (Lundberg *et al.*, 2011). Briefly, binding of paired cDNA-tagged antibodies directed against the targeted serum proteins lead to hybridization of the corresponding DNA oligonucleotides allowing subsequent extension by a DNA polymerase. The protein level is quantified using real-time PCR. Only samples that passed the quality control tests are reported. If expression was below the detection limit, the value reported is the lower limit of detection. Only proteins that were detectable in at least 50% of samples were used for subsequent analysis.

### Statistical analysis

The statistical analysis was performed using GraphPad Prism (version 8.4.3, GraphPad Software, La Jolla California USA) and R software (version 4.0.1) using the package “mgcv”. Mann-Whitney Wilcoxon test was used to test for differences between continuous variables and p-values were adjusted for multiple testing using the Holm method. Categorical variables were compared using Fisher’s exact test. Generalized additive models were used to evaluate relationships between time since symptom onset and different variables, with the number of knots used to represent the smooth term set at three.

The correlations between cellular subsets and the serum protein expressions were analyzed using non-parametric Spearman correlations. The significance threshold was set at an alpha < 0.05. For the differential expression analysis, a false-discovery rate (Benjamini, Krieger and Yekutieli, 2006) of 1% was used as significance threshold, except for the late vs early comparison where 5% was used as indicated.

## Author contribution

SC, YZ and CC contributed to study design, patient recruitment, data collection, data analysis, data interpretation, and writing of the manuscript. SS, AJ and SC developed the CyTOF antibody panel and performed the CyTOF experiments. SA contributed to experiments, data collection, data analysis, data interpretation. MER, EB, AR, MS-H, LCH, ASZ, and DJS contributed to patient recruitment and clinical management. NDS contributed to writing of the manuscript. JN, OB and BB contributed to study conception and design, data analysis, data interpretation, and writing of the manuscript. All authors reviewed and approved the final version of the manuscript.

## Acknowledgments

We thank Alessandra Guaita, Jennifer Jörger, Sara Hasler and the members of the Bodenmiller and Boyman laboratories for their support and helpful discussions.

