## Supplementary material for "A distinct innate immune signature marks progression from mild to severe COVID-19"

### Supplementary Figure Legends

**Figure S1: Experimental design for mass cytometry data acquisition in two independent batches and quality assessment. Related to Figure 1.**

(A) (A) Schematic of the strategy used to collect mass cytometry data. Samples were analyzed in two batches. The indicated number of samples were barcoded and stained with the same antibody mix stored as frozen aliquots.

(B) Histograms of signal intensities for each marker included in the panel observed on the pooled references acquired in the two runs. Data were linearly scaled based on the 98$^{th}$ percentile. Shifts in background intensity were observed for CD16, CD66ace, and Granzyme B.

(C) t-SNE plot calculated for the three reference samples included in each CyTOF run based on all markers included in the study, colored by batch

(D) Dotplot of signal intensities for CD16 versus CD15 in reference sample.

(E) Manual gating strategy used to identify the main cell types used to train the random forest cell classifier.

(F) t-SNE plot as in Figure 1C of a random subset of 1000 immune cells from each sample colored by normalized marker intensity.

**Figure S2: In-depth characterization of the neutrophil and monocyte compartments. Related to Figure 2**

(A) t-SNE plots as in Figure 2A of normalized expression of the indicated markers across a maximum of 1000 neutrophils per patient colored by clusters identified with the PhenoGraph algorithm.

(B) Heatmap of the normalized marker expression for the PhenoGraph neutrophil clusters. Each cluster was manually assigned to three neutrophil subsets based on the negative, intermediate, or high expression of CD16.

(C) Visualization of monocytes clusters using first and second components of a diffusion map. Cells are colored by PhenoGraph clusters. The two branches leading to the CD169+ cells are shown with black arrows.

(D) Diffusion map as described in A colored by marker

**Figure S3: In-depth characterization of the neutrophil and monocyte compartments. Related to Figure 3.**

(A) Heatmap of correlations between monocyte and neutrophil clusters and myeloid immune subsets across the cohort. The colors indicate Pearson's correlation coefficients.

**Figure S4: Disease status of patients with high cluster 4 frequency, related to Figure 4.**

(A) Scatter plot of M4 cluster frequencies relative to the time after symptom onset, including hospitalization status. The cluster frequency is given in relation to the total monocytes in the PBMCs.

**Figure S5: Association between serum protein levels, disease stages, and myeloid subsets. Related to Figure 5.**

(A) Scatter plots of expression of selected cytokines as measured by ELISA. The dots are colored by disease grade at sampling time. The expression levels of the healthy controls are shown as a reference on the left. The pseudo-time course was modeled using a general additive model for the disease severities separately (mild, blue lines; severe, red lines).

(B) Scatter plots of the expression of the indicated serum proteins versus the time after symptom onset. The expression is given as NPX on a log2 scale. The dots are colored by disease grade at sampling time. The expression levels in healthy controls are shown as a reference on the left. The pseudo-time course was modeled using a general additive model for the disease severities separately (mild, blue lines; severe, red lines).

(C) Volcano plot of proteomics data for COVID-19 patients with mild vs. severe disease at early (top panel) and late (bottom panel) stages. An FDR of 5% was taken as significance cut-off.

(D) Heatmap of Spearman correlation coefficients for relationships between the different PhenoGraph clusters, myeloid immune cell subsets, and significant differentially expressed serum proteins.

(E) Scatter plots of frequencies of the indicated clusters versus expression of selected serum proteins in individual patients as described in Figure 5E, but colored by time after symptom onset.

(F) Scatter plots of frequencies of the indicated clusters versus expression of selected serum proteins in individual patients as described in Figure 5E, but colored by time after symptom onset.

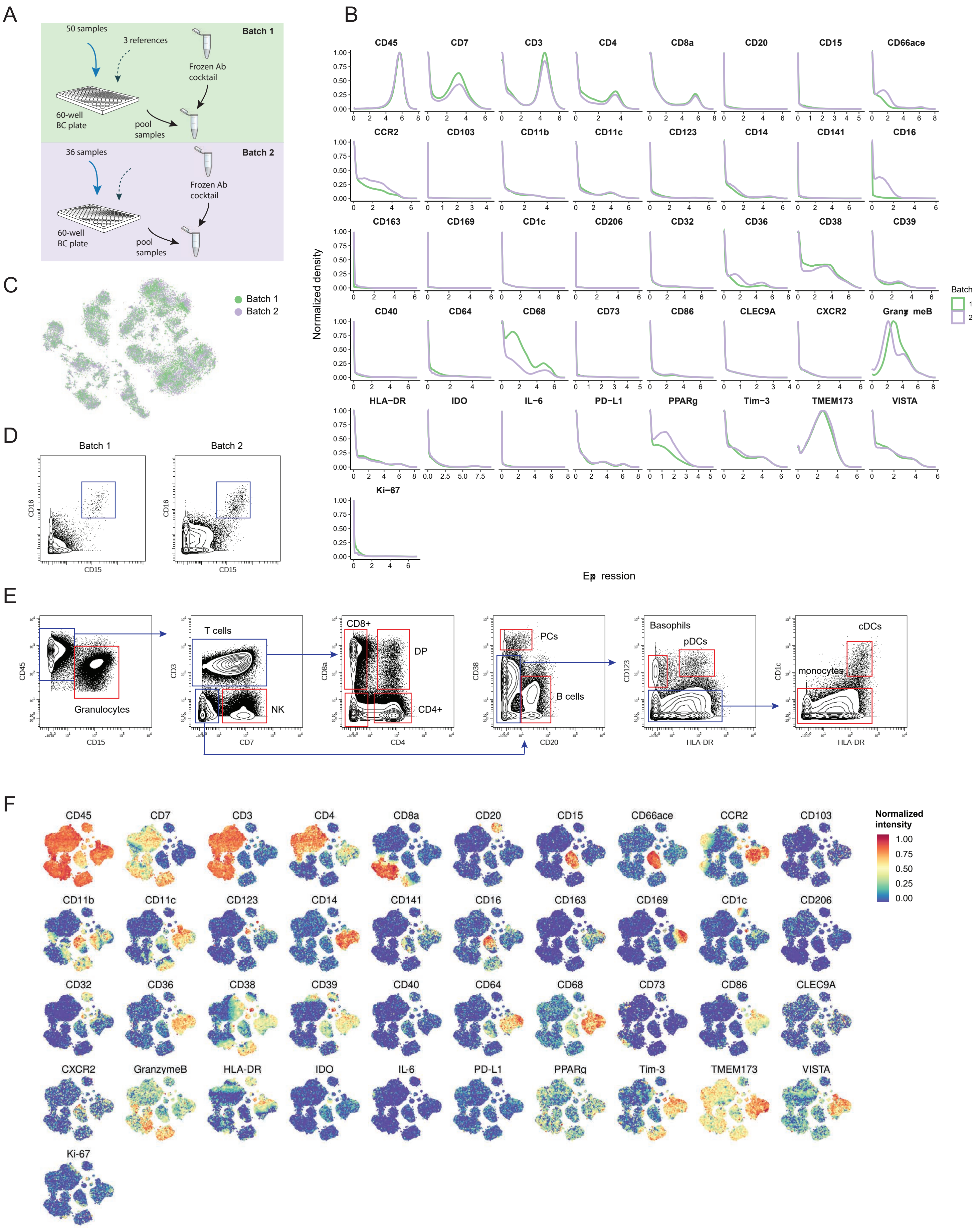


**Figure S1.** Experimental design for mass cytometry data acquisition in two independent batches and quality assessment. Related to Figure 1.

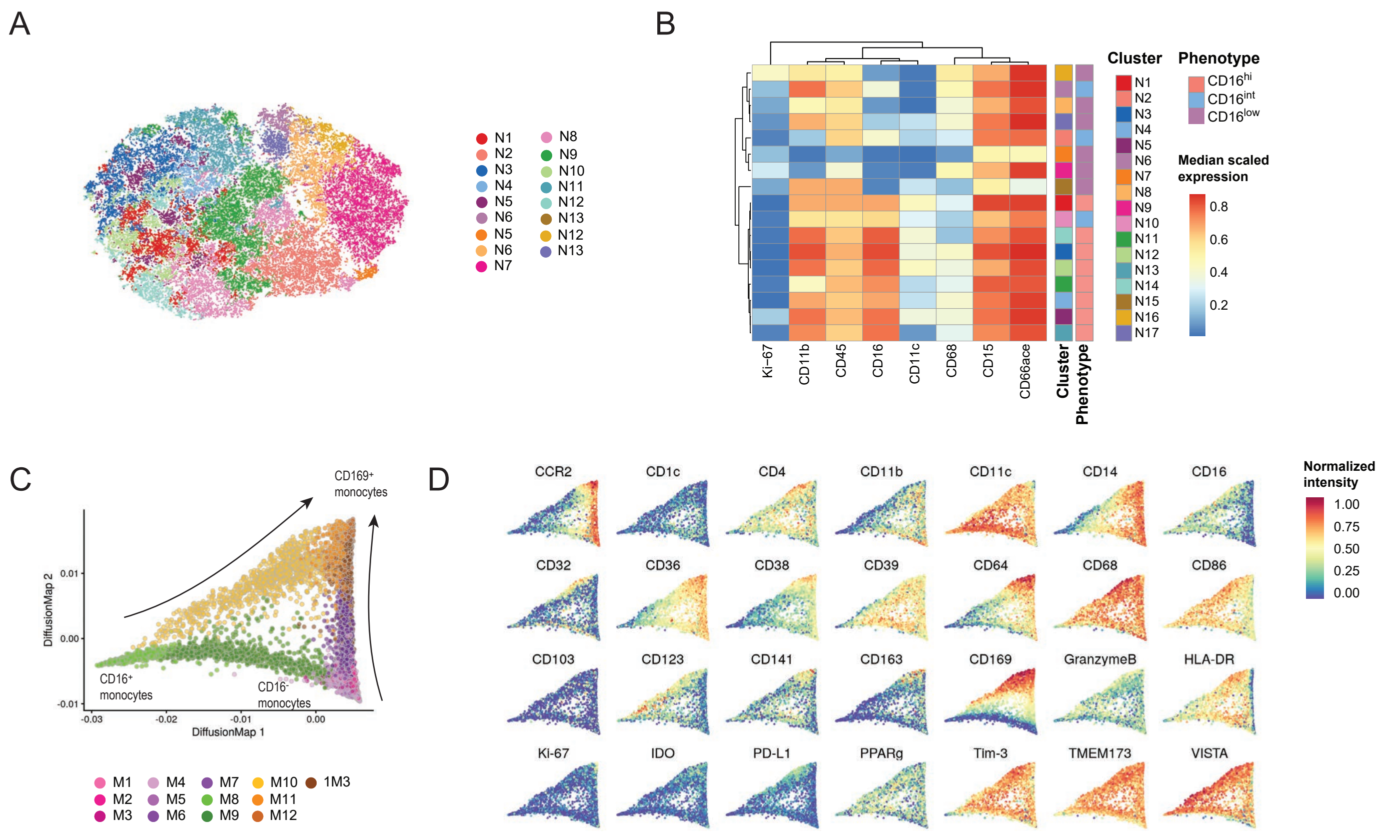


**Figure S2.** In depth characterization of the neutrophil and monocyte compartment. Related to Figure 2.

A

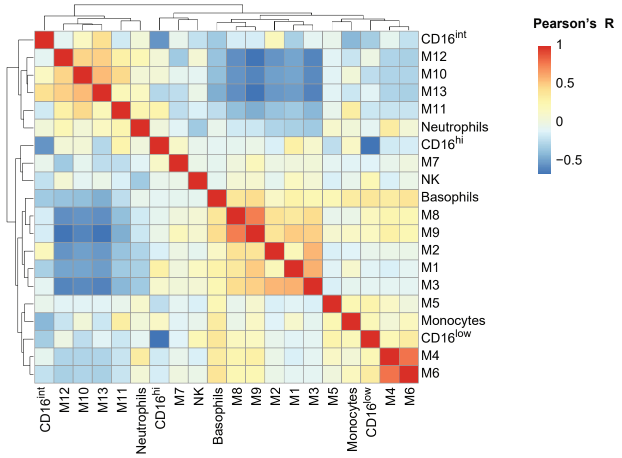


**Figure S3.** Correlation between myeloid subsets and other immune components. Related to Figure 3.

A

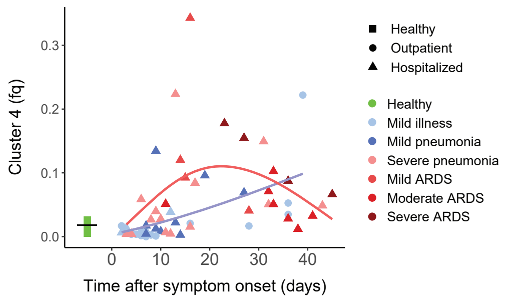


**Figure S4.** Patient care at inclusion timepoint of patients with high cluster 4 frequency. Related to Figure 4.

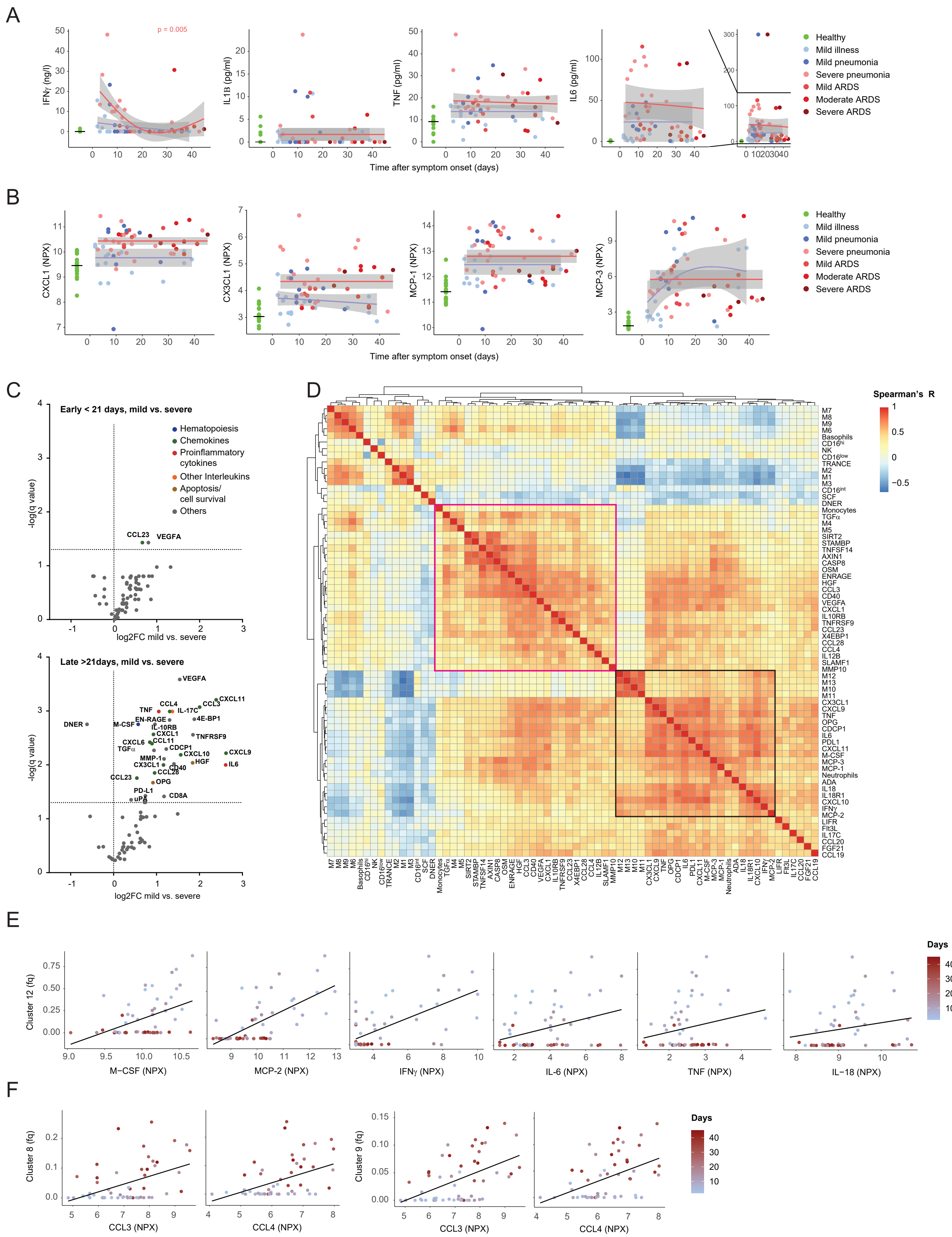

**Figure S5.** Association between serum protein levels, disease stages and myeloid subsets. Related to Figure 5.

### Table S1: Markers used for Mass Cytometry Analysis

| Antigen | Description | Provider | Clone | Cat No | Metal tag |
|---|---|---|---|---|---|
| **Cell lineage** | | | | | |
| CD45 | Pan immune | BioLegend | HI30 | 304002 | 209Bi |
| CD1c | Dendritic cells | BioLegend | L161 | 331502 | 172Yb |
| CD3 | T cells | BioLegend | UCHT1 | 300402 | 195Pt |
| CD4 | CD4 T cells | BioLegend | RPA-T4 | 300502 | 145Nd |
| CD7 | NK cells | BD Bioscience | M-T701 | 555359 | 196Pt |
| CD8a | CD8 T cells | BioLegend | RPA-T8 | 301002 | 175Lu |
| CD15 | Granulocytes | Biolegend | HI98 | 301902 | 89Y |
| CD20 | B cells | BD Bioscience | H1(FB1) | 555677 | 148Nd |
| CD66a/c/e | Granulocytes | BioLegend | ASL-32 | 342302 | 161Dy |
| **Fc & complement receptors** | | | | | |
| CD11b | Complement R. 3 | BioLegend | M1/70 | 101202 | 147Sm |
| CD16 | Low affinity FCGR3a | BioLegend | 3G8 | 302002 | 158Gd |
| CD32 | Low affinity FCGR2a | BioLegend | Fun-2 | 303202 | 156Gd |
| CD64 | High affinity FCGR1a | BioLegend | 10.1 | 305002 | 141Pr |
| **TLR/CLR/Cytokine receptors** | | | | | |
| CD14 | LPS co-Receptor | BioLegend | M5E2 | 301802 | 168Er |
| CD123 | IL-3 Receptor | BioLegend | 6H6 | 306002 | 150Nd |
| CD370 | C-type lectin receptor | Abcam | EPR22324 | ab245121 | 144Nd |
| **Ectoenzymes** | | | | | |
| CD38 | cADP ribose hydrolase | BioLegend | HIT2 | 303502 | 149Sm |
| CD39 | Ectonucleotidase | BioLegend | A1 | 328202 | 151Eu |
| CD73 | Ectonucleotidase | BioLegend | AD2 | 344002 | 155Gd |
| **Cell adhesion** | | | | | |
| CD11c | Integrin | BioLegend | Bu15 | 337202 | 174Yb |
| CD68 | Glycoprotein | eBioscience | KP1 | 14-0688-82 | 143Nd |
| CD103 | Integrin | Abcam | SP301 | ab245746 | 166Er |
| CD169 | Sialoadhesin | BioLegend | 7-239 | 346002 | 163Dy |
| CD206 | Mannose receptor | BioLegend | 15-2 | 321102 | 146Nd |
| **Migration** | | | | | |
| CCR2 | C-C chemokine R. 2 | BioLegend | K036C2 | 357202 | 159Tb |
| CXCR2 | C-X-C chemokine R. 2 | R&D Systems | 48311 | MAB331-100 | 142Nd |
| **Immunomodulatory Molecules** | | | | | |
| TIM-3 | Coinhibitory R. | R&D Systems | Polyclonal | AF2365 | 167Er |
| VISTA | Coinhibitory R. | Fluidigm | D1L2G | 3160025D | 160Gd |
| CD274 (PD-L1) | Coinhibitory R. | CST | E1L3N | 13684BF | 154Sm |
| CD40 | Costimulatory R. | eBioscience | 5c3 | 14-0409-82 | 152Sm |
| CD86 | Costimulatory R. | BD Bioscience | 2331 (FUN-1) | 555655 | 171Yb |
| CD141 | Thrombomodulin | BioLegend | M80 | 344102 | 162Dy |
| Indoleamine 2,3-dioxygenase | Immunomodulatory mediator | Abcam | SP260 | ab245737 | 164Dy |

| | | | | | |
|---|---|---|---|---|---|
| IL-6 | Proinflammatory cytokine | BioLegend | MQ-13A5 | 501115 | 153Eu |
| Granzyme B | Cytotoxic Protease | Invitrogen Antibodies | GB11 | MA1-80734 | 173Yb |
| HLA-DR | Ag presentation | BioLegend | L243 | 307602 | 194Pt |
| TMEM173 | Stimulator of interferon genes | Abcam | SP339 | ab238796 | 170Er |
| **Scavanger receptors** | | | | | |
| CD36 | Class B S.R. | BioLegend | 5-271 | 336202 | 165Ho |
| CD163 | High affinity S.R. | BioLegend | GHI/61 | 333602 | 169Tm |
| **Transcription factors** | | | | | |
| PPARg | Nuclear receptor | BioLegend | 14G4B17 | 683402 | 176Yb |
| **Cell division** | | | | | |
| Ki-67 | Proliferation marker | BD Bioscience | B56 | 556003 | 198Pt |
